# Site-specific phosphorylation of tau impacts mitochondrial biology and response to stressors

**DOI:** 10.1101/2023.02.19.529131

**Authors:** Michael O Isei, Peter A Girardi, Joel Rodwell-Bullock, Keith Nehrke, Gail VW Johnson

## Abstract

Phosphorylation of tau at sites associated with Alzheimer’s disease (AD) likely plays a role in the disease progression. Mitochondrial impairment, correlating with increased presence of phosphorylated tau, has been identified as a contributing factor to neurodegenerative processes in AD. However, how tau phosphorylated at specific sites impacts mitochondrial function has not been fully defined. We examined how AD-relevant phosphomimetics of tau impact selected aspects of mitochondrial biology. To mimic phosphorylation at AD-associated sites, the Ser/Thr sites in wild-type GFP tagged-tau (T4) were converted to glutamic acid (E) to make pseudophosphorylated GFP tagged-Ser-396/404 (2EC) and GFP tagged-Thr-231/Ser-235 (2EM) constructs. These constructs were expressed in neuronal HT22 cells and their impact on specific mitochondrial functions and responses to stressors were measured. Phosphomimetic tau altered mitochondrial distribution. Specifically, mitochondria accumulated in the soma of cells expressing either 2EC or 2EM, and neurite-like extensions in 2EC cells were shorter. Additionally, ATP levels were reduced in both 2EC and 2EM expressing cells, and ROS production increased in 2EC cells during oxidation of succinate when compared to T4 expressing cells. Thapsigargin reduced mitochondrial membrane potential (Ψ_m_) and increased ROS production in both 2EC and 2EM cells relative to T4 cells, with no significant difference in the effects of rotenone. These results show that tau phosphorylation at specific AD-relevant epitopes negatively affects mitochondria, with the extent of dysfunction and stress response varying according to the sites of phosphorylation. Altogether, these findings extend our understanding of potential mechanisms whereby phosphorylated tau promotes mitochondria dysfunction in tauopathies, including AD.

**Funding information:** R01 AG067617

## Introduction

Tau is a unique phosphoprotein predominantly expressed in the neurons (Weingarten et al., 1975; Paterno et al., 2022). Functionally, it was originally associated with microtubule stabilization and assembly, however it is now evident that tau is involved in numerous cellular processes including mitochondria-trafficking and signaling, synaptic activity and gene expression (Ittner et al., 2008; Tracy et al., 2022). Tau functions are regulated by extensive posttranslational modifications, especially phosphorylation. Its longest isoform (2N4R) comprises 441 amino acids and 85 potential serine (Ser), threonine (Thr) and tyrosine (Tyr) sites where phosphorylation can occur (Xia et al., 2021). While modification at some sites is not implicated in disease, aberrant phosphorylation at specific sites can convert tau to a pathological protein (Alonso et al., 2010). Indeed, specific soluble and oligomeric species of tau that are abnormally phosphorylated at AD associated sites have been identified as pathogenic forms (Zheng et al., 2020; Xia et al., 2021). Accumulation of pathogenic tau is highly associated with neuronal damage and cognitive defects observed in AD (Xia et al., 2021; Trease et al., 2022).

Immunostaining with tau antibodies and mass spectrometry have aided the identification of specific sites in tau that are predominantly phosphorylated in AD brain (Kimura et al., 2018; Wesseling et al., 2020). Phosphorylation at Thr-181 and Thr-231 occur early in AD and likely play a key role in facilitating subsequent phosphorylation events (Wesseling et al., 2020; Stefanoska et al., 2022). Tau phosphorylated at Thr-205, Ser-396 and Ser-404 has been shown to be associated with neuronal impairment in AD (Mondragón-Rodríguez et al., 2014; Xia et al., 2021). In addition, the abundance of tau phosphorylated at Ser-202/Thr-205 has been extensively studied in AD and specific antibodies targeting these sites have been developed to monitor AD progression (Neddens et al., 2018). Despite our understanding of the contribution of aberrantly phosphorylated tau in AD pathogenesis, its mechanisms of toxicity remain to be fully elucidated. This results, in part, from our partial understanding of the impacts of abnormally phosphorylated tau on other proteins and organelles.

In addition to accumulation of phosphorylated tau, impaired mitochondrial function is an early event in the evolution of AD (Reddy, 2011; Eckert et al., 2014; Grimm et al., 2017). This indicates that there may be an interplay between abnormal tau phosphorylation and mitochondria dysfunction. Reports of tau localization in different mitochondrial compartments and its interaction with mitochondrial proteins further underscore the relationship between tau and mitochondria (Drummond et al., 2020; Tracy et al., 2022). Tau is also involved in mediating neuronal mitochondria transport (Pérez et al., 2018). Thus, it is not unexpected that tau phosphorylated at AD-relevant sites can directly impact mitochondrial functions. However, information on the impact of AD-relevant tau phosphorylation on mitochondria biology in basal and stressed conditions is not extensive. Previous investigations have revealed that tau phosphorylated at Ser-396/404 accumulates in the mitochondria of presynaptic neurons in aged mice, thereby increasing ROS generation (Torres et al., 2021, 2022). Phosphomimetic at Thr-231 impairs stress-induced mitophagy in *C. elegans* (Guha et al., 2020), whereas mitochondrial distribution is disrupted in PC12 cells and mouse neurons expressing tau phosphorylated at Ser-202 and Thr-205 (Shahpasand et al., 2012). Although these findings suggest that mitochondrial functions can be directly impacted by pathogenic tau, further investigations into how different AD-relevant tau phosphorylation influence mitochondrial functions are needed. Such studies will enhance our understanding of the mechanisms whereby different tau modifications induce toxicity in AD pathology.

In this study, a mouse hippocampal cell line (HT22) transiently expressing wild type GFP-tagged tau (T4) as well as phosphomimetic GFP-tagged tau at Ser-396/404 (2EC) or Thr-231/Ser235 (2EM) were used to examine the effects of site-specific tau phosphorylation on mitochondrial functional parameters. Our results revealed that phosphomimetic tau impairs mitochondria distribution, reduces ATP production, and increases ROS generation. We demonstrate that tau phosphorylated at AD-relevant sites impairs cellular response to stressors, particularly endoplasmic reticulum-derived calcium stress.

## Methods and Materials

### Reagents

Dulbecco’s modified Eagle medium (DMEM, #2418283), gentamicin (#2328211) and GlutaMax (#2380959) were from GIBCO, Thermo Fisher Scientific (MA, USA). Glucose (#50997) was from Alfa Aesar, Thermo Fisher Scientific (MA, USA). Thapsigargin (#10522) and carbonyl cyanide 4- (trifluoromethoxy) phenylhydrazone (FCCP, #15218) were purchased from Cayman chemical company (MI, USA). Rotenone (#150154) was from MP Biomedicals (OH, USA). Tetramethylrhodamine methyl ester (TMRM, #2321830), horseradish peroxidase (HRP, #9003990) and AmplexUltra-Red (#2365697) were from Invitrogen Thermo Fisher Scientific (OR, USA). Fetal clone II (FCII, #AF29578030) was from Cytiva Hyclone (UT, USA) while paraformaldehyde (#15710) was from Electron Microscopy Sciences (PA, USA). L-malic acid (#97676), L-glutamic acid (#56860) and L-succinic acid (#110156) were from MilliporeSigma (MA, USA).

### Cell culture and constructs

HT22 cells were provided by Dr. P.H Reddy, Texas Tech University. The cells were cultured in DMEM supplemented with 10% fetal clone II (FCII), 2.2 mM GlutaMax, 25 µg/ml gentamicin and maintained at 37 °C in a 5% CO_2_ incubator. Mitochondrial targeted mCherry was from Addgene (#55102). GFP-tagged human wild type tau (0N4R: GFP-T4) and its pseudophosphorylated (serine and/or threonine changed to glutamic acid) forms (GFP-T42EM: T231E/S235E (2EM) and GFP-T42EC: S396E/S404E (2EC)) were used. GFP-T4 and GFP-T42EC have been described previously (Quintanilla et al., 2014). GFP-T42EM was created by cloning T42EM (Ding et al., 2006) into the HindIII and BamHI sites of the pEGFP-C1 plasmid (6084-1, Clontech). To verify tau protein expression, cells grown until 50-60% confluent on 60 mm dishes were transfected with 0.375 µg empty vector (EV), wild type or phosphomimetic tau plasmids using PolyJet (SL100688, SignaGen Laboratories) according to manufacturer’s instructions. Protein expression was confirmed using western blot analysis.

### Immunoblotting

Cells were gently washed with phosphate buffer saline (PBS) on ice and lysed with RIPA buffer containing protease inhibitors. Thereafter, the lysates were sonicated (Misonix Inc, S-3000) for 10 s, then centrifuged at 4 °C for 10 min at 16 000g. The resulting supernatants were collected, and protein concentration was measured using BCA assay. Protein was then denatured in 5x SDS buffer for 10 min at 100 °C. Equal protein amounts (20 µg) were loaded and separated on a 12% SDS-PAGE gel and transferred onto a nitrocellulose membrane. The membrane was blocked for 60 min at room temperature with 5% non-fat milk prepared in Tris-buffered saline containing 0.05% Tween 20 (TBS-T). Then, appropriate primary antibody, rabbit anti-tau (1:40000, A0024, Dako), was added and incubated overnight at 4 °C. After washing three times with TBS-T at 15 min intervals, the nitrocellulose membranes were incubated with appropriate HRP-conjugated secondary antibody, anti-rabbit HRP (1:5000, R1006, KwikQuant, Kindle Biosciences) in 5% non-fat milk for 60 min at room temperature. Protein bands were revealed with enhanced chemiluminescence reagent (211595, Millipore, Immobilon Crescendo) and images captured with a KwikQuant Imager (Kindle Biosciences). The nitrocellulose membranes were subsequently stripped and re-probed with rabbit anti-GAPDH (1:5000, sc-25778, Santa Cruz Biotechnology, Inc.) for protein loading control.

### Effect of phosphomimetic tau on mitochondrial distribution

To assess the impacts of phosphomimetic tau on mitochondrial distribution, cells were grown on glass coverslips coated with poly-d-lysine (PDL; Sigma Aldrich, A003E). The cells were allowed to grow until 50-60% confluent prior transfection with 0.75 µg total cDNA [0.375 µg mitochondrial targeted-mCherry (mCherry) and 0.375 µg of GFP-tagged EV, T4, 2EM or 2EC]. The transfection media was replaced with freshly warmed media 16 h post-transfection then incubated for 48 h. Thereafter, the media was aspirated and cells were gently washed three times with pre-warmed PBS while slightly rocking on a shaker for 5 min. The cells were then fixed with a freshly prepared 4% paraformaldehyde-sucrose solution for 5 min and washed three times at 5 min interval with PBS. The coverslips were mounted on glass-microscope slides using Fluoro-Gel Mounting Medium (Electron Microscopy Sciences, 17985-30). Representative pictures were taken with Zeiss microscope equipped with Colibri 7 LED and Axiocam 705 camera, using 40x oil immersion objective. Excitation for GFP and mCherry were 488 nm and 590 nm, respectively, with emission at 509 nm and 610 nm. Mitochondrial distribution and abundance were assessed using Fiji Image J software.

### Determination of mitochondrial membrane potential and ATP levels

Changes in mitochondrial membrane potential (ΔΨ_m_) were determined using TMRM, a cell-permeable cationic dye that accumulates in the mitochondria. We used an optimized non-quench concentration (20 nM) of TMRM that generates fluorescence as a direct function of Ψ_m_. Cells grown on 25 mm PDL-coated glass coverslips were gently washed twice with PBS-glucose buffer and incubated with TMRM prepared in PBS-glucose buffer for 20 min. Thereafter, the coverslips were encased in an Attofluor™ Cell Chamber (ThermoFisher, Cat# A7816) and mounted on the microscope stage. Baseline TMRM fluorescence was recorded for 5 min prior addition of 10 µM FCCP to uncouple the mitochondria depolarize Ψ_m_. To assess the impact of stressors on ΔΨ_m_, the cells were treated with 1 µM thapsigargin (a non-competitive inhibitor of sarcoendoplasmic reticulum Ca^2+^-ATPase) or 1 µM rotenone (a mitochondrial complex I ubiquinone binding site inhibitor) after baseline TMRM fluorescence was established. Notably, the cells were maintained in TMRM in PBS-glucose buffer during imaging to prevent re-equilibration of the dye across the mitochondria membrane (Debattisti et al., 2017; Connolly et al., 2018). Imaging was performed with Zeiss microscope equipped with Colibri 7 LED and Axiocam 705 camera under constant acquisition conditions across all experiment, including, resolution, time-interval and -cycle. In addition, the laser power and intensity were reduced to 5% to avoid photobleaching. Changes in TMRM fluorescence intensity were determined using 40x oil immersion objective following excitation and emission at 548 nm and 573 nm, respectively. About five to ten transfected cells per image were analyzed using Fiji Image J software to determine the ΔΨ_m_. Background fluorescence was subtracted and ΔΨ_m_ was expressed as the ratio between the baseline fluorescence intensity and change in fluorescence intensity after stimuli (F_o_/F_o_-F_1_). F_o_ is the mean baseline fluorescence intensity while F_1_ is the mean fluorescence intensity upon exposure to stressor (Santos et al., 2005; Quintanilla et al., 2013). Data were normalized with the total number of mitochondria per cell.

ATP levels were measured using an ATP determination Kit (Invitrogen, #A22066) and read on a Glomax Luminometer (Promega). Briefly, cells were grown in 24-well plates at a seeding density of 2 × 10^4^ cells/ml. Cells were collected 48 h post-transfection in lysis buffer containing 100 mM Tris-HCl, 4 mM EDTA, 0.1% Triton X-100, and protease inhibitor. Thereafter, the cells were sonicated and centrifuged to remove cell debris. Standards curve of known ATP concentration (0.031 - 2 µM) was generated and used to calibrate ATP concentration in samples. Data obtained were normalized to protein concentration measured using BCA assay.

### Effect of phosphomimetic tau on ROS production

ROS generation was monitored using AmplexUltra-Red-HRP-hydrogen peroxide (H_2_O_2_) detection system. Briefly, cells were plated at 5 × 10^4^ cells per well in 48-well plates and were transfected accordingly at 50-60% confluent. Forty-eight hours post-transfection, the cells were gently washed with PBS-glucose buffer and permeabilized for 2 min with 5 µg/ml saponin in PBS-glucose buffer. Thereafter, the cells were incubated with 5 U/ml HRP and substrate (5 mM succinate or 5 mM malate plus 2 mM glutamate) for 5 min followed by addition of Amplex-UltraRed (50 μM final well concentration) for a 200 μl final assay volume. In this reaction, HRP catalyzes H_2_O_2_-oxidation of AmplexUltra-Red to resorufin-like AmplexUltrox-Red that emits fluorescence at 590 nm upon excitation at 530 nm. We measured kinetic data for 30 min at 90 s intervals with Multi-mode microplate reader (Biotech Synergy LX, 22011326). For each experiment, standard curve generated with freshly prepared H_2_O_2_ (0 - 5 μM) was used to calibrate fluorescence intensity to pmoles units of H_2_O_2_. We subtracted background fluorescence measured in wells containing PBS-glucose only. To assess the effect of stressors on ROS production, the cells were incubated with rotenone (1 μM) or thapsigargin (1 μM) for 10 min prior energization with substrates.

### Statistics

Statistics were performed with GraphPad Prism 9 (GraphPad Software, Inc.) and analysis were carried out using one-way ANOVA followed by Tukey’s post hoc test. Data are expressed as means ± SEM and statistical differences were expressed as p < 0.05 *, p < 0.005 **, p < 0.001 ***, p < 0.0001 ****.

## Results

### Expression of phosphomimetic tau altered mitochondrial distribution and abundance

Prior to initiating our studies, we first confirmed that all the tau constructs were expressed at equivalent levels. The immunoblot in Fig. 1A shows that all the GFP-tagged tau constructs expressed at similar levels in HT22 cells. The slight differences in electrophoretic mobility of 2EC and 2EM protein compared with T4 correlate with the modifications at the respective Ser/Thr sites (Ding et al., 2006).

**Fig. 1.**
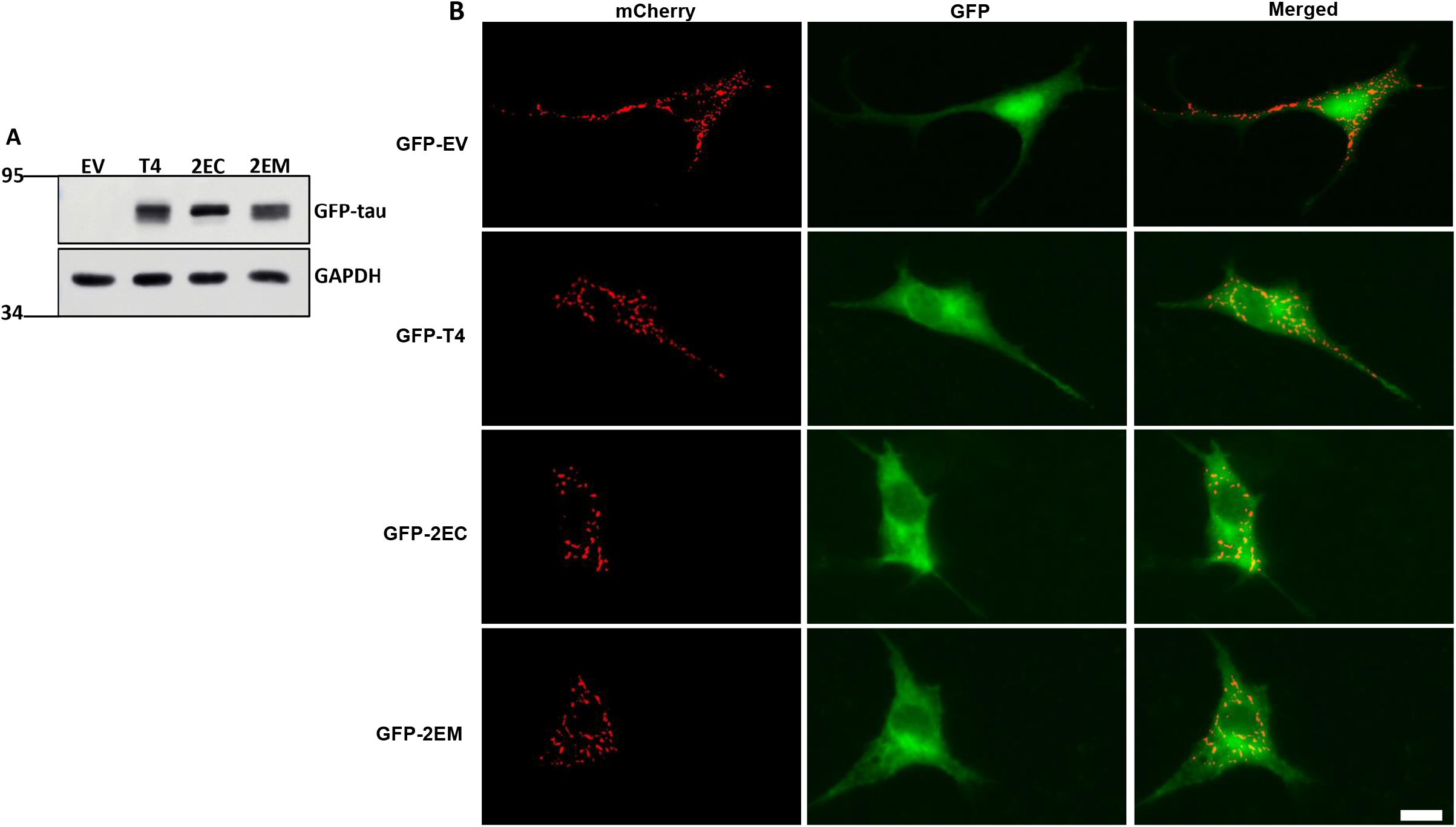
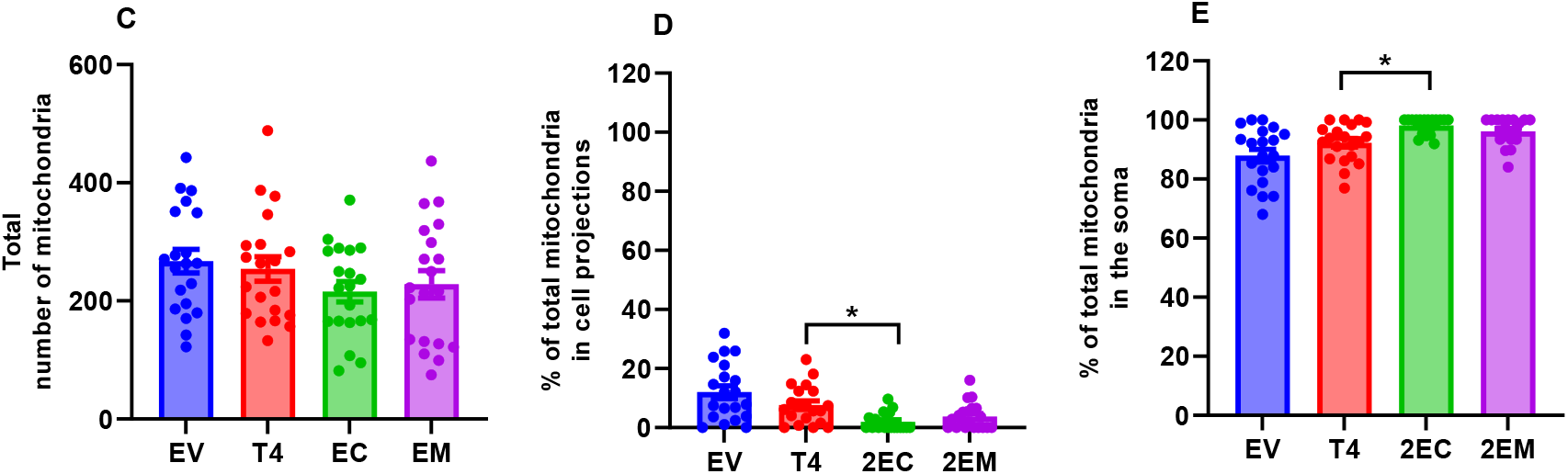
Tau expression and effect of phosphomimetic tau on mitochondrial distribution and abundance. (A) Representative immunoblot for total tau in cell lysates (20 µg protein). GFP tagged wild-type tau (T4) and phosphomimetic tau (2EC and 2EM) are equivalently expressed in HT22 cells. The empty vector (EV) lane indicates the absence or very low levels of endogenous tau in HT22 cells. The cell lysates were probed with a tau antibody and GAPDH was used as loading control. HT22 cells were co-transfected with mitochondrial targeted mCherry and EV or the indicated GFP-tau constructs prior to fixation and imaging. Representative images of merged, mito-mCherry mitochondria-puncta and GFP showing shorter cell projections in cells expressing 2EC and 2EM. Scale bar is 20 µm (B). The total number of mitochondria per cell (C), % of total mitochondria in the projections (D) and soma (E) were normalized to fluorescence intensity. Quantification was performed on twenty cells from four independent experiments. Data analysis was by one-way ANOVA followed by Tukey’s multiple-comparisons test and presented as mean ± SEM. p < 0.05 *.

Neuronal cells depend on appropriate mitochondrial distribution to maintain function and neural connections. Aberrant distribution alters synaptic activities while inadequate neuronal-mitochondrial populations contribute to neuronal damage observed in AD (Harerimana et al., 2022). To determine if the presence of abnormally phosphorylated tau could impact mitochondrial distribution and population, cells were transfected with mito-mCherry and GFP tagged-T4 or the different phosphomimetic forms of tau. Fig. 1B shows that cells expressing phosphomimetic tau appear to have shorter projections compared to T4 and EV, with a significant accumulation of mitochondria in the soma. Thus, there was a significant reduction in percent of total mitochondria in the projections in cells expressing 2EC compared to T4 (Fig. 1D,E).

### Impact of phosphomimetic tau on mitochondrial membrane potential (Ψ_m_)

Ψ_m_ is a common measure of mitochondrial function and reflects the charge or electrical component of the proton motive force that drives oxidative phosphorylation. Hence, we asked whether phosphorylated tau impacts Ψ_m._ To this end, we measured Ψ_m_ using TMRM.

Treatment with the mitochondrial membrane uncoupler FCCP significantly decreased the TMRM fluorescence intensity, verifying that the fluorescence was indeed originating from the mitochondria. Quantitative representation of changes in Ψ_m_ indicates no significant difference in all tau constructs (Fig. 2A,B). Next, we examined the impact of specific stressors on ΔΨ_m_. Rotenone suppressed Ψ_m_, but the observed changes were not significantly different across of groups (Fig. 3A,B). However, compared to T4, inhibition of SERCA with thapsigargin significantly decreased Ψ_m_ in cells expressing both 2EC and 2EM (Fig. 3C,D), suggesting that cells expressing phosphomimetic tau were more susceptible to calcium stress. Further, ATP levels in cells expressing 2EC and 2EM phosphomimetic tau were significantly reduced compared to T4 expressing cells (Fig. 4) suggesting decreased oxidative phosphorylation.

**Fig. 2.**
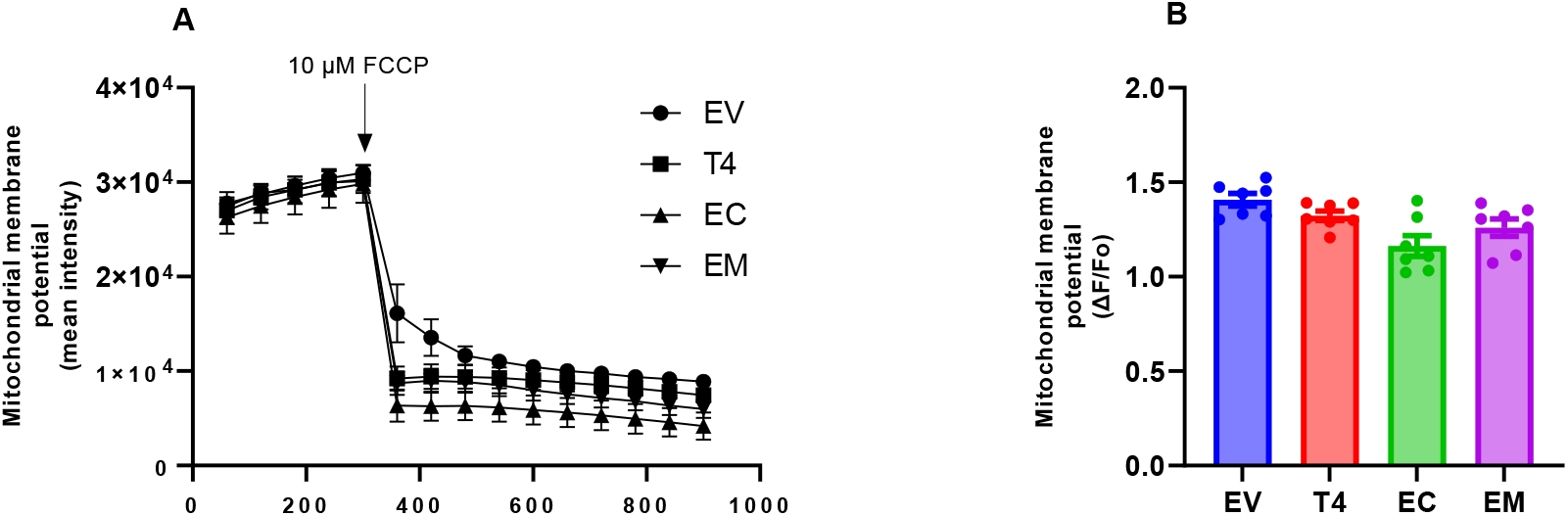
Assessment of mitochondrial membrane potential in cells expressing wild type and phosphomimetic tau. Tracings showing ΔΨ_m_ in cells expressing GFP-empty vector (EV), or T4, 2EC and 2EM, respectively. After taking the baseline TMRM fluorescence for 5 min, the cells were treated with FCCP, and the fluorescence changes were recorded for another 10 min at 1 min interval (A). The decreased TMRM fluorescence intensity upon addition of FCCP indicates reduced Ψ_m_ and that the dye signal was from the mitochondria. Quantitative representation of the baseline TMRM fluorescence intensity after 5 min normalized with mitochondrial abundance (B). Each data point represents the mean TMRM fluorescence intensity of 5 to 10 transfected cells in seven independent experiments. Data analysis was by one-way ANOVA followed by Tukey’s multiple-comparisons test.

**Fig. 3.**
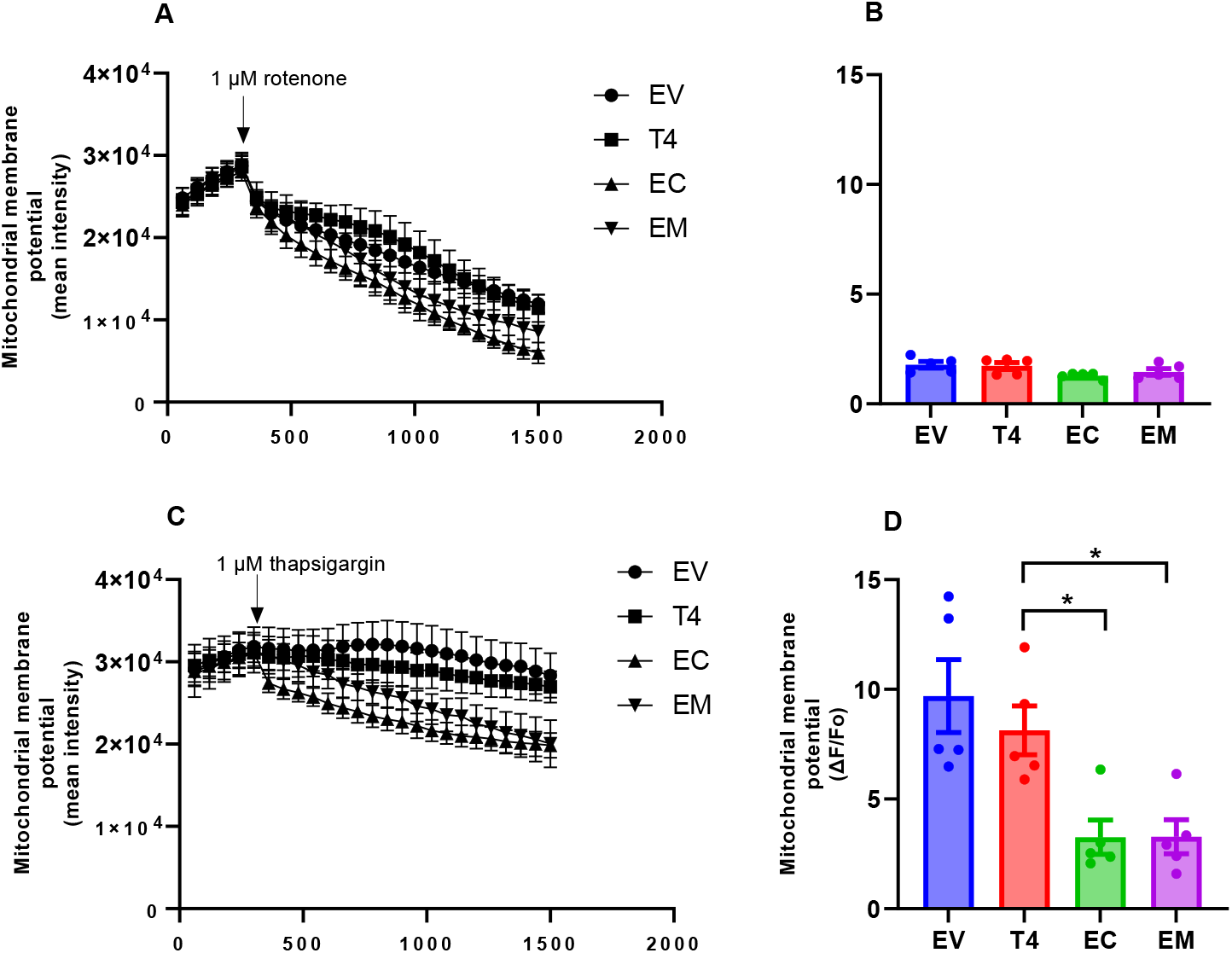
Stressors reduced mitochondrial membrane potential cells expressing phosphomimetic tau. Transfected cells were treated with 1 µM rotenone or 1 µM thapsigargin, respectively, after establishing the baseline TMRM fluorescence for 5 min. Representative tracings and quantitative representation of ΔΨ_m_, respectively, following exposure to rotenone (A,B) or thapsigargin (C,D). Changes in TMRM fluorescence intensity were recorded for 20 min at 1 min interval upon treatment with stressors. After subtracting background fluorescence, fluorescence intensities were estimated using the formula ΔF/F_o_ where ΔF represents changes in TMRM fluorescence intensity before treatment, while F_o_ is the difference after exposure to stressor. Fluorescence intensity was normalized with mitochondria abundance. Each data point represents the mean fluorescence intensity of 5 to 10 transfected cells in five independent experiments. Data analysis was by one-way ANOVA followed by Tukey’s multiple-comparisons test and presented as mean ± SEM. p < 0.05 *.

**Fig. 4.**
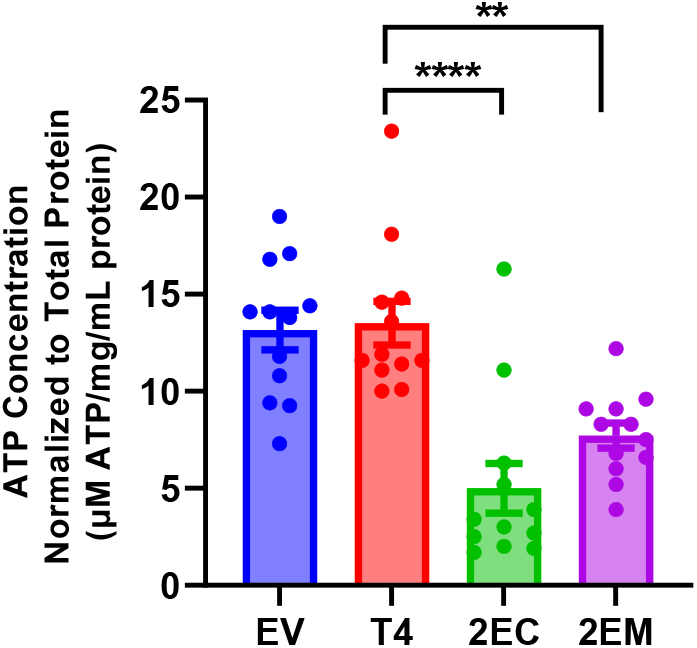
Phosphomimetic tau altered ATP production. ATP level was assessed using an ATP determination kit according to the manufacturer’s instructions. Expression of phosphomimetic tau significantly reduced ATP levels. Data analysis was by one-way ANOVA followed by Tukey’s multiple-comparisons test and presented as mean ± SEM. p < 0.005 **, p < 0.001 ****. n = 3.

### Phosphomimetic tau altered ROS production

Mitochondria are the primary site of ROS formation which in pathological conditions can result in oxidative stress. Because aberrantly phosphorylated tau has been suggested to increase oxidative stress and thus contribute to AD progression (Torres et al., 2021), we tested whether tau pseudophosphorylated at AD-associated sites altered ROS formation. We measured ROS production in unenergized cells, during oxidation of different metabolic substrates and upon exposure to stressors. ROS generated by unenergized mitochondria and during oxidation of malate-glutamate was not significantly different across all groups (Fig. 5A,B). However, compared to T4, ROS production increased significantly in 2EC expressing cells during oxidation of succinate (Fig. 5C). Irrespective of the tau construct, ROS production was the highest during oxidation of succinate. Therefore, we assessed the impact of rotenone or thapsigargin on ROS production during oxidation of succinate. Our data indicated that rotenone did not alter ROS generation across all groups (Fig. 5D). In contrast, thapsigargin significantly increased ROS generation in all groups (Fig. 5E). Calculating the difference between stressed and unstressed states (Fig 5F,G) showed that ROS generation in the presence of thapsigargin was significantly greater in 2EC and 2EM-expressing cells compared to T4 (Fig. 5G).

**Fig. 5.**
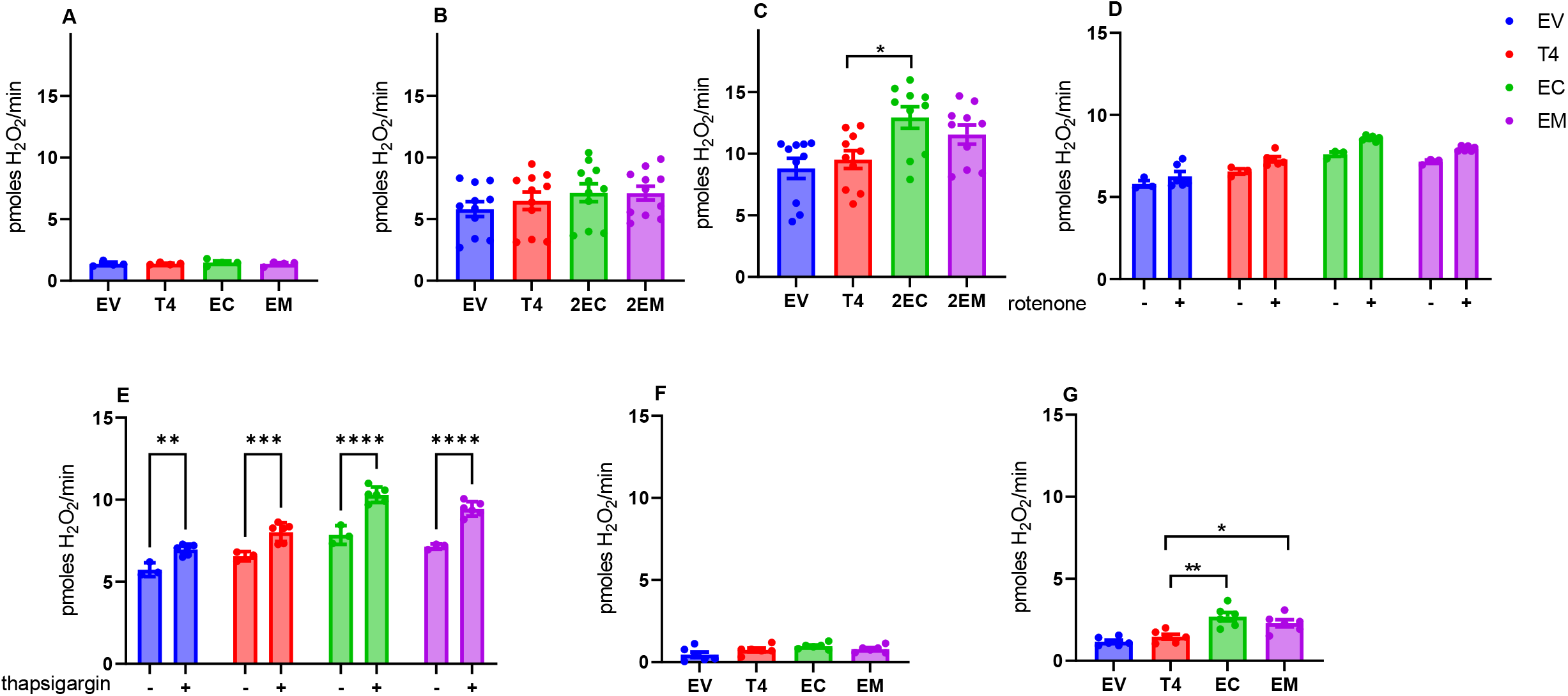
Effect of phosphomimetic tau on ROS generation. ROS production by unenergized mitochondria (A) and during oxidation of 5 µM malate-2 µM glutamate (B), or 5 µM succinate (C). Changes in ROS generation by mitochondria oxidizing succinate following exposure to rotenone (D,F) or thapsigargin (E,G). Data in F and G represent the difference in ROS generation between the unstressed state (C) and after treatment with rotenone (F) or thapsigargin (G). Data analysis was by one-way ANOVA followed by Tukey’s multiple-comparisons test and presented as mean ± SEM. p < 0.05 *, p < 0.005 **. n = 3.

## Discussion

Previous reports have demonstrated that phosphorylated tau induces neuronal damage by disrupting microtubule assembly and stabilization (Johnson and Stoothoff, 2004; Iqbal et al., 2008; Di et al., 2015). However, emerging evidence suggests that the damage caused by phosphorylated tau may transcend microtubule-associated effects, due to its interaction with other cellular components, including the mitochondria (Szabo et al., 2020; Tracy et al., 2022). Since healthy mitochondria are essential for neuronal survival, understanding the impact of pathogenic forms of tau on mitochondrial functions has become a key area of focus in AD pathology. Our data demonstrate that tau pseudophosphorylated at AD-associated sites alters mitochondrial functional parameters, with the magnitude of the impact depending on the sites where tau is phosphorylated. Our findings extend understanding of how phosphorylated tau causes mitochondrial dysfunction and support the possible mechanistic rationale for mitochondrial-oriented therapies in AD.

We have demonstrated that phosphorylated tau impairs mitochondrial distribution. Our finding is consistent with previous reports in PC12 cells expressing phosphomimetics tau at Ser-202 and Thr-205 (Shahpasand et al., 2012) and in P301L tau knock-in mouse neurons (Rodríguez-Martín et al., 2016). Specifically, a higher percentage of mitochondria puncta accumulated in the soma, in a manner similar to findings in cellular and mouse models of AD (Biernat and Mandelkow, 1999; Kopeikina et al., 2011; Shahpasand et al., 2012). Mitochondrial accumulation in the soma and corresponding decrease in neurite-like extensions may be attributed to the defective neurite outgrowth we observed in cells expressing phosphomimetic tau. We observed identical results when the cells were stained for tubulin (Fig. S4). Indeed, 2EC and 2EM mutations impair microtubule assembly and stabilization that is essential for neurite growth (Johnson and Stoothoff, 2004; Ding et al., 2006) thereby disrupting mitochondria transport and resulting in accumulation in the soma. Impaired mitochondria transport and accumulation in neuronal soma are important arbiters of synaptic dysfunction in AD (Wang et al., 2015; Cheng and Bai, 2018). Of note, a significantly greater percentage of total mitochondria puncta binned at 5 – 10 µm^2^ were found in cells expressing 2EC compared with T4 (Fig. S1-3), suggesting an impairment in mitochondria dynamics (Szabo et al., 2020). Further investigation is needed to probe the mechanism by which tau phosphorylation affects mitochondrial dynamics and evaluate the potential implications for neurodegenerative diseases such as AD.

In physiological state, neuronal energy supply is achieved largely by effective mitochondrial bioenergetics, which include metabolic substrate catabolism, electron transfer along the mitochondrial electron transport system (mETS), and generation of Ψ_m_. Ψ_m_ is crucial for ATP synthesis via oxidative phosphorylation, and its assessment can be used as a proxy to reflect mitochondrial energetic state in AD (Eckert et al., 2014; Pérez et al., 2018; Szabo et al., 2020). Basal Ψ_m_ was not different across all tau constructs when normalized with mitochondria abundance, suggesting that phosphorylated tau does not significantly influence mechanisms involved in regulating Ψ_m_ in the absence of stressors. However, ATP levels were reduced in cells expressing 2EC and 2EM, indicating that phosphomimetic tau might impair oxidative phosphorylation. Our findings underscore studies in SH-SY5Y cells and transgenic mice expressing P301L tau (David et al., 2005; Schulz et al., 2012) and a report (Torres et al., 2022) that phosphorylated tau accumulates in synaptic mitochondria resulting in decreased ATP production. Interestingly, Drummond et al. (2020) and Torres et al. (2021, 2022) reveal that tau phosphorylated at Ser-396/404 (2EC) interacts with the β and O subunit (ATP5B and ATP5O) of complex V, which has been associated with impaired mitochondrial bioenergetics, including decreased ATP synthesis (Rhein et al., 2009; Torres et al., 2022). While proteomic analysis is required to identify potential downstream targets of 2EM and 2EC, it is conceivable that both tau phosphomimetics interact with specific mitochondrial proteins to disrupt ATP synthesis.

High oxygen consumption rate predisposes the brain to elevated ROS production (Cobley et al., 2018), and ROS generation in AD brain can be intensified by the presence of hyperphosphorylated tau (Rhein et al., 2009; Eckert et al., 2014). ROS emission during oxidation of succinate increased in cells expressing phosphomimetic tau compared to T4. Succinate oxidation stimulates ROS generation from complex III through forward electron transfer and/or by reverse electron transport (RET) to complex I which is sensitive to inhibition by rotenone (Brand, 2016). We found that ROS generation was insensitive to rotenone suggesting the absence of RET in HT22 cells and that phosphomimetic tau plausibly stimulates ROS production at sites downstream of complex I. Additionally, increased ROS production likely results from accumulation of defective mitochondria, which could be due in part to phosphorylated tau-induced defective mitophagy (Reddy and Oliver, 2019; Guha et al., 2020). Intracellular calcium plays fundamental regulatory and signaling roles in neuronal functions and its concentration is tightly regulated by the endoplasmic reticulum (ER) and mitochondria (Bagur and Hajnóczky, 2017). Disturbances in calcium homeostasis have been linked to tau-mediated toxicity in AD (Berridge, 2013; Palikaras et al., 2021). Thapsigargin, a classical ER-calcium stressor, precipitates intracellular calcium overload by blocking SERCA-mediated calcium transport from the cytoplasm into the ER (Sehgal et al., 2017). Thapsigargin-induced ER-calcium stress decreased Ψ_m_ and increased ROS generation in cells expressing 2EC and 2EM consistent with our previous findings in immortalized cortical neurons expressing Asp-421-truncated tau (Quintanilla et al., 2009) and immortalized striatal progenitor cells expressing mutant huntingtin (STHdh^Q111/Q111^) (Quintanilla et al., 2013). These findings provide evidence that tau phosphorylated at specific AD-relevant sites impairs cellular response to calcium stress and that ER stress-induced calcium dysregulation may be a key mechanism of phosphorylated tau-mediated neurodegeneration in AD.

Even with extensive research effort, there is no detailed understanding of the specific cellular changes that link abnormally phosphorylated tau with the progression of AD. In this study, we demonstrated that tau phosphorylated at AD-relevant sites precipitate several indicators of mitochondrial dysfunction reported in human AD patients, including impaired stress response, decreased ATP levels, and increased oxidative stress. Our results support the association between aberrantly phosphorylated tau and defective mitochondria in the pathogenesis of AD (Szabo et al., 2020; Torres et al., 2022). Our findings suggest that changes in mitochondria function can be used to integrate the mechanism of toxicity of different AD-relevant tau phosphorylation species. Additionally, our observations are based on a tau overexpression model, leaving open the possibility that in physiological or low expression levels, tau may impose a different, yet undetermined mitochondria effect. Further research is needed to determine the likelihood of a synergistic adverse effect of abnormal tau phosphorylation and mitochondria dysfunction in the progression of AD.

## Abbreviations

2EC: GFP tagged-Ser-396/404
2EM: GFP tagged-Thr-231/Ser-235
2N4R: two N-terminal and four repeat domains
AD: Alzheimer’s disease
ATP: adenosine triphosphate
EV: empty vector
FCCP: carbonyl cyanide 4- (trifluoromethoxy) phenylhydrazone
GFP: green fluorescent protein
HRP: horseradish peroxidase
HT22 cells: Immortalized mouse hippocampal neuronal cell line
PBS: phosphate buffer saline
PDL: poly-d-lysine
ROS: reactive oxygen species
Ser: Serine
T4: GFP tagged-tau
Thr: threonine
TMRM: Tetramethylrhodamine methyl ester
Tyr: tyrosine
Ψ_m_: mitochondrial membrane potential

## Acknowledgements

We would like to thank Jacen Emerson for his assistance, and Dr. P.H Reddy for his generous gift of HT22 cells. This work was supported by the National Institutes of Health (NIH) R01 AG067617.

## Authors contributions

GVWJ and KN contributed to the study conception and design, edited the manuscript, and provided funding. MOI contributed to the study conception and design, performed experiments, analyzed the data, interpreted the experiments and wrote the manuscript. PG contributed to the study conception and design, performed experiments, analyzed the data. JR-B performed experiments, analyzed the data.

## Supplemental Figures

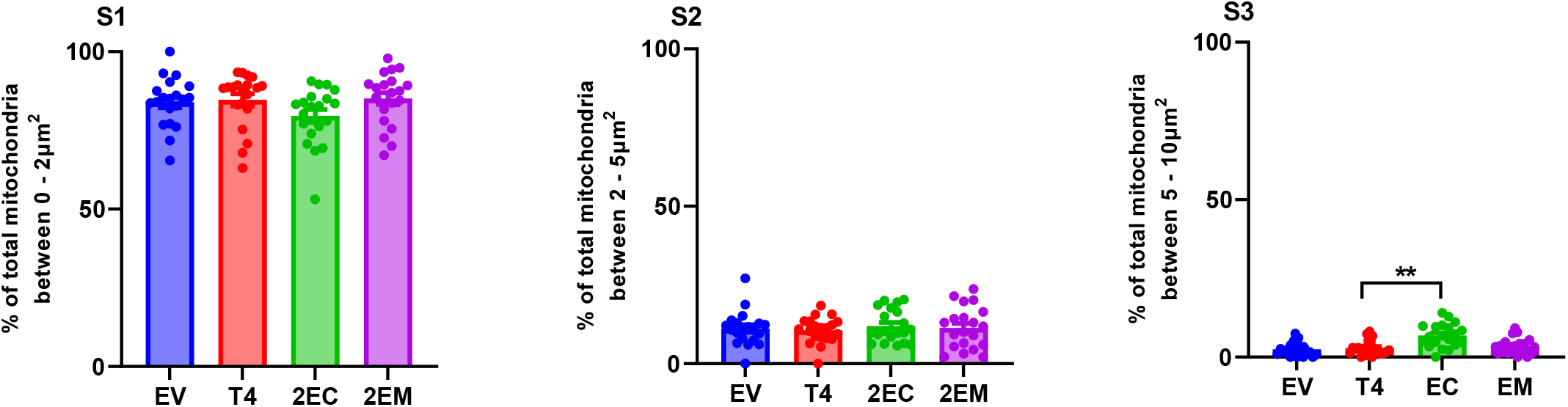

**Effect of phosphomimetic tau on mitochondrial abundance**

% of total mitochondria binned between 0-2 µm^2^ **(S1)**, 2-5 µm^2^ **(S2)**, and 5-10 µm^2^ **(S3)**. Quantification was performed on twenty cells from four independent experiments. Data analysis was by one-way ANOVA followed by Tukey’s multiple-comparisons test and presented as mean ± SEM. **p < 0.005**.

**Cells expressing phosphomimetic tau have short neurite-like extensions (S4)** Images showing shorter neurite-like extensions in cells expressing 2EC and 2EM relative to T4. Scale bar is 20 µm Transfected cells were gently washed twice with PBS, then fixed with 4 % paraformaldehyde for 10 min at room temperature. Thereafter, the cells were washed three times with PBS and permeabilized with 0.25% Trixton-X-100 in PBS for 10 minutes. The cells were blocked with 5% BSA and 300 mM glycine for 60 min, then labelled overnight with primary antibody (beta tubulin #10094-1, Proteintech, 1:200), followed by two rinses in PBS, and incubation with secondary antibodies (Alexa Fluor 594, ThermoFisher Scientific, 1:1000) for 60 min at room temperature. The coverslips were then mounted on glass slides with Fluoro-Gel Mounting Medium and observed using a Zeiss microscope equipped with Colibri 7 LED and Axiocam 705 camera, using a 40x oil immersion objective.

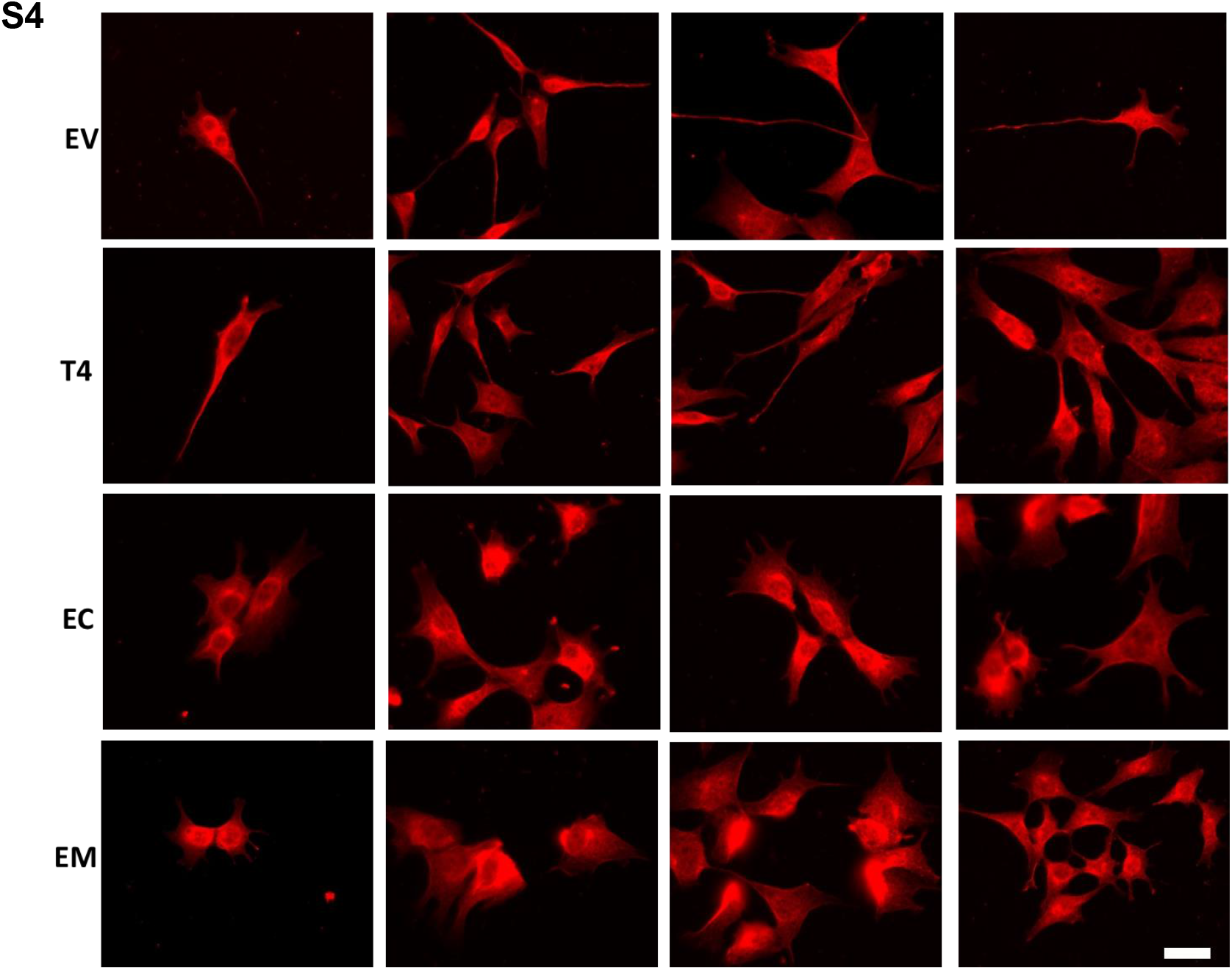

